# Individual differences in the influence of mental imagery on conscious perception

**DOI:** 10.1101/607770

**Authors:** N. Dijkstra, M. Hinne, S.E. Bosch, M.A.J. van Gerven

## Abstract

Mental imagery and visual perception rely on similar neural mechanisms, but the function of this overlap remains unclear. One idea is that imagery can influence perception. Previous research has shown that imagining a stimulus prior to binocular presentation of rivalling stimuli increases the chance of perceiving the imagined stimulus. In this study we investigated how this effect interacts with bottom-up sensory input by comparing psychometric response curves for congruent and incongruent imagery in humans. A Bayesian hierarchical model was used, allowing us to simultaneously study group-level effects as well as effects for individual participants. We found strong effects of both imagery as well as its interaction with sensory evidence within individual participants. However, the direction of these effects were highly variable between individuals, leading to weak effects at the group level. This highlights the heterogeneity of conscious perception and emphasizes the need for individualized investigation of such complex cognitive processes.

---

In daily life, we are bombarded with visual stimuli. When walking down the street, we see different colours, shapes and textures. At the same time, we are often caught up in thinking about future or past events, which is accompanied by mental imagery (Delamillieure et al., 2010). Mental imagery and visual perception rely on similar neural mechanisms (Dijkstra, Bosch, & van Gerven, 2019). Throughout the visual system, imagining objects and perceiving them elicits similar activation patterns (Dijkstra, Bosch, & van Gerven, 2017; Lee, Kravitz, & Baker, 2012; Reddy, Tsuchiya, & Serre, 2010) and similar top-down connectivity (Dijkstra, Zeidman, Ondobaka, Van Gerven, & Friston, 2017). Given that we often engage in mental imagery while receiving visual input, the question arises to what extent imagery influences perception.

Few studies have directly explored the interaction between imagery and perception. One line of work has focused on the behavioural effects of imagery on conscious perception during binocular rivalry. Binocular rivalry is a phenomenon in which a different image is presented to each eye of an observer. In general, the observer will only perceive one of the images consciously, while the other image is suppressed (Levelt, 1966). Pearson and colleagues (2008) showed that imagining one of two stimuli prior to binocular rivalry leads to a priming effect: it biases perception towards the imagined stimulus. In contrast, *perceiving* one of the two stimuli prior to binocular rivalry leads to adaptation: decreasing the chance of subsequently perceiving that stimulus (Knapen, Brascamp, van Ee, Kanai, & van den Berg, 2007). The priming effect of imagery has been dissociated from effects of attention (Pearson, Clifford, & Tong, 2008) and is specific to the orientation (Pearson et al., 2008) and location of the stimulus (Chang, Lewis, & Pearson, 2013). This means that imagery can influence conscious perception.

However, the exact nature of this influence remains unclear. According to predictive coding theories of perception, top-down signals should interact with bottom-up sensory evidence (Bastos et al., 2012; den Ouden, Kok, & de Lange, 2012; Friston, 2005). In the case of binocular rivalry, sensory evidence can be defined as the relative contrast of the two stimuli: increasing the contrast of one stimulus while keeping the other constant increases the proportion of trials where that stimulus is consciously perceived (Carter & Cavanagh, 2007; Chen & He, 2004; Levelt, 1966). This can be illustrated clearly with a psychometric curve (see Fig. 1A). In this study, we investigated how top-down imagery and bottom-up sensory input interact by characterizing the effect of imagery on the parameters of this psychometric curve.

**Figure 1.**
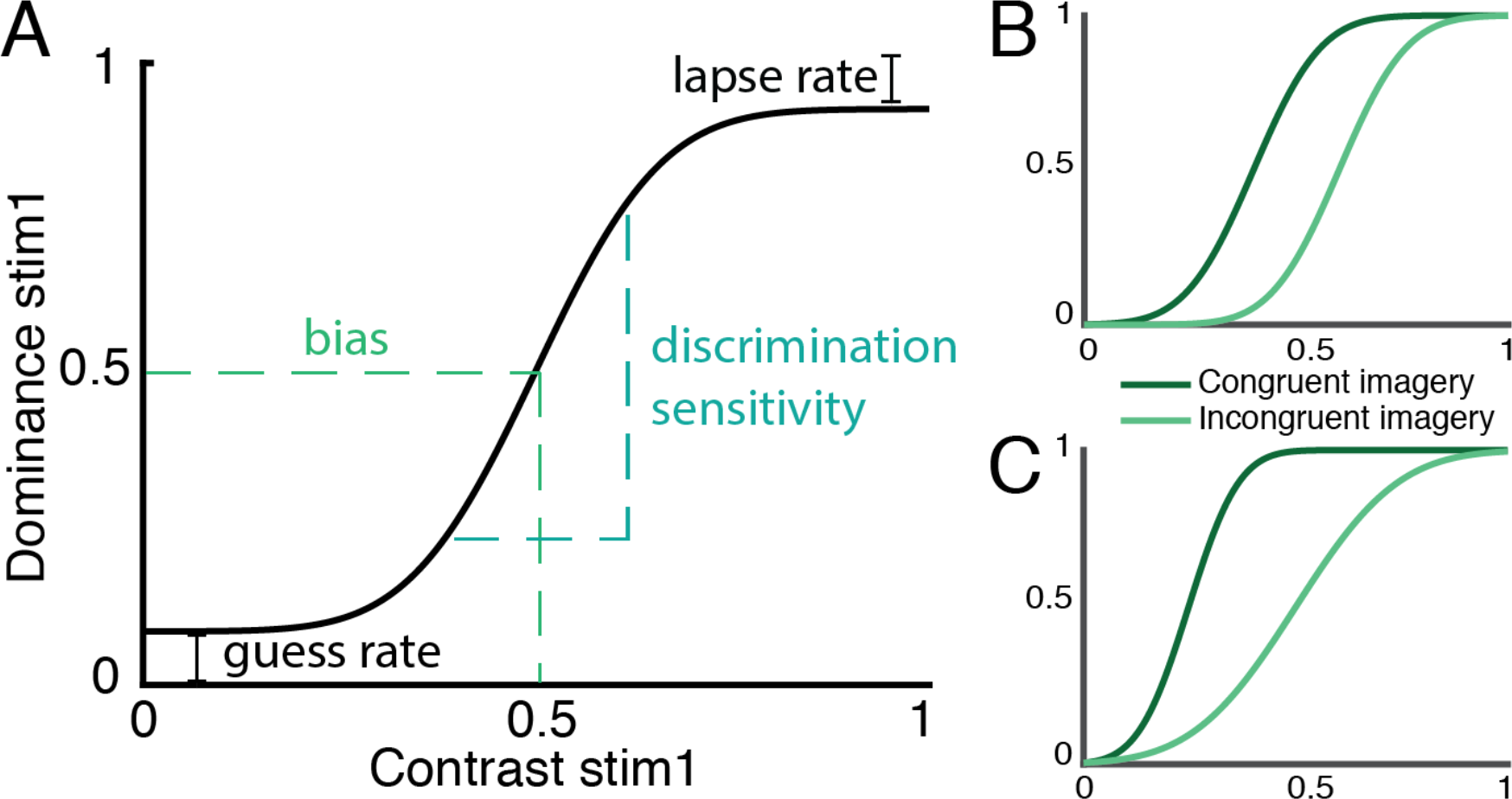
Illustration of psychometric curves. The contrast of the manipulated stimulus is plotted on the x-axis. The contrast of the other stimulus is fixed at 0.4. The y-axis indicates the proportion of trials on which the manipulated stimulus is reported dominant. (A) Standard psychometric curve based on the algorithm by Wichmann & Hill (2001). Four parameters determine the shape of the curve. The *bias* is the required contrast value to reach a dominance of 0.5. The *discrimination sensitivity* or slope is the difference in contrast value between 0.25 and 0.75 dominance. The *guess rate* is the dominance associated with a contrast of 0 and the *lapse rate* is 1 - the dominance associated with a contrast of 1. (B) Main effect of imagery is reflected in a shift in the bias during congruent versus incongruent imagery. (C) Interaction with sensory evidence is reflected in a change in discrimination sensitivity during congruent versus incongruent imagery.

One possibility is that there is no interaction, but that imagery and sensory evidence influence perception independently. This would lead to a constant shift in the psychometric curve, such that for all dominance levels, a lower contrast is needed to achieve that dominance level when imagining the congruent stimulus compared to the incongruent stimulus (Fig. 1B). This would be reflected in a lower *bias* or offset for congruent versus incongruent imagery (Wichmann & Hill, 2001). Alternatively, top-down imagery may interact in a systematic way with bottom-up sensory input. This would mean that the difference between congruent and incongruent imagery varies at different dominance levels. In terms of the psychometric curve, this would result in an effect of imagery on the steepness of the curve, which is a measure of the *discrimination sensitivity* (Fig. 1C). Imagining a stimulus then makes people more or less sensitive to changes in congruent bottom-up sensory input.

The nature and strength of mental imagery varies greatly between individuals (Galton, 1880; Kozhevnikov, Kosslyn, & Shephard, 2005; Marks, 1973) and between clinical populations (Matthews, Collins, Thakkar, & Park, 2014; Sack, van de Ven, Etschenberg, Schatz, & Linden, 2005; Shine et al., 2014). Therefore, we also investigated whether there are individual differences in the interaction between imagery and perception. Determining whether different types of interactions exist is important for the direction of future research and could have clinical implications.

We used an adapted version of the binocular rivalry imagery task developed by Pearson et al. (2008). The sensory evidence of the binocular rivalry display was manipulated by changing the relative contrast of the stimuli. A hierarchical Bayesian model was used to fit psychometric curves per participant on the dominance levels during congruent and incongruent imagery. We then compared the derived parameters between the conditions. This approach allowed us to characterize the presence as well as the absence of effects on the group level and on the individual participant level.

## Materials and Methods

### Participants

Sixty-one participants gave written informed consent and participated in the study. One participant was excluded due to a very high response bias (above two standard deviations from the mean) and one was excluded because no responses were logged due to technical issues. Fifty-nine participants were included in the reported analyses (mean age ± *SD* = 23.10 ± 2.59). The study was approved by the local ethics committee (Commissie Mensgebonden Onderzoek region Arnhem-Nijmegen). To determine the number of participants required, we performed a power calculation (see *Supplementary Material 1.1*).

### Apparatus

The experiment took place in a darkened room with dark walls. Participants sat with their head on a chin rest, and wore red-blue anaglyph glasses. The stimuli were generated using MATLAB and the Psychophysics Toolbox (RRID: rid_000041) and presented on a luminance-calibrated Iiyama Vision Master CRT screen (resolution: 1280 × 1024 pixels, refresh rate: 86 Hz). The distance to the screen was 42.5 cm. Stimuli comprised coloured sinusoidal annular gratings of 5.56 degrees of visual angle (red stimulus oriented at 45°, blue stimulus oriented at 135°) that were presented around a central fixation bulls-eye (radius, 0.5°). The luminance-value of the red stimulus at full contrast was set to half the luminance-value of the blue stimulus.

### Experimental paradigm

We adapted the imagery binocular rivalry task from Bergmann et al. (2015). Prior to the experimental task, participants filled in the Vividness of Visual Imagery Questionnaire 2 (VVIQ2) (Marks, 1973; Marks, 1995). Then, eye-dominance was determined (see *Manipulating sensory evidence*) and participants’ binocular stereovision was tested. After that, the rivalry stimulus was shown continuously for a brief time to familiarise participants with the binocular stimulus. Participants briefly practised the main task, and were given the opportunity to ask questions.

The task started with a fixation bulls-eye, after which a cue indicated whether to imagine the red (‘R’) or blue (‘B’) grating (Fig. 2). Participants imagined the cued grating for 7 s and then indicated their experienced imagery vividness using a sliding bar on a scale from −150 to 150 where values below 0 indicated low vividness and values above 0 indicated high vividness. If no response was given within 3 s, the trial continued. Subsequently, the rivalry stimulus was presented for 750 ms, after which participants indicated whether they saw the red stimulus, a perfect mixture or the blue stimulus. Participants were instructed to only indicate ‘mixed’ if both stimuli were equally dominant. Again, if no response was given within three seconds, the experiment continued. Reponses were recorded with a keyboard. Vividness ratings were given with the left hand and rivalry responses were given with the right hand.

**Figure 2.**
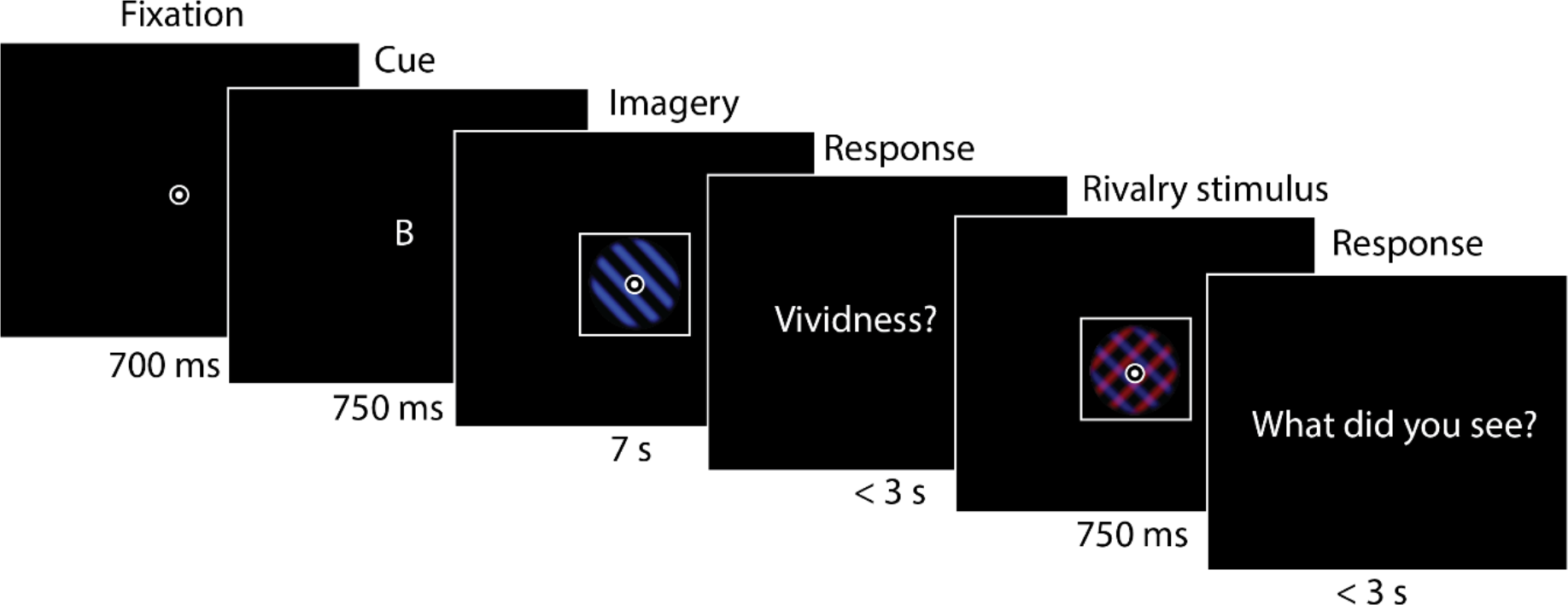
Experimental design. After fixation, a cue was presented for 750 ms indicating whether participants had to imagine the blue grating (‘B’) or the red grating (‘R’). The cued grating was then imagined as vividly as possible for 7 s, after which participants indicated their experienced imagery vividness on a continuous scale using a sliding bar. Next, the rivalry stimulus was presented for 750 ms. Participants indicated whether they saw the blue grating, the red grating or a perfect mixture of the two.

To control for response bias, ten percent of the trials were catch trials. These trials
contained mock-rivalry stimuli which were a spatial mix of half a red and half a blue stimulus. These displays had blurred edges and the exact split varied on each trial to resemble actual piecemeal rivalry (Keogh & Pearson, 2017; Pearson, Rademaker, & Tong, 2011). On these trials, participants should respond ‘mixed’ and there should be no effect of imagery on the response for these trials. If there was an effect of imagery, this indicated that the participant had a response bias.

### Manipulating sensory evidence

We varied the contrast of one grating between 0 and 1 and kept the contrast of the other grating fixed at 0.4. To be able to test our hypotheses, we wanted to estimate the full psychometric curve. Low dominance levels are easily obtained when the contrast of the manipulated grating is set to zero. However, high dominance levels are more difficult to obtain, because even when the manipulated grating has a contrast value of 1, the fixed grating with a contrast of 0.4 could still dominate in a large percentage of the trials if the participant has a strong bias towards the eye corresponding to the fixed grating. Therefore, we decided to manipulate the grating corresponding to the dominant eye of the participant. This increased the chance of reaching high dominance levels for the manipulated grating. We determined which eye was dominant using a variation of the Miles test (Miles, 1930; Roth, Lora, & Heilman, 2002). During this test, participants view an object through a triangle created by holding their hands together, thumbs extended and arms outstretched. They focus on the object with both of their eyes open and then close one eye. If the object shifts outside of the triangle, that eye is the dominant eye.

### Deriving dominance per contrast

For each participant, and for each condition, we measured 100 contrast levels between 0 and 1. To infer the dominance levels at different contrast values, we calculated the percentage the manipulated grating was dominant using a sliding window over 11 subsequent contrast values. Mixed responses were counted as a dominance of 0.5. For instance, for the 11 trials with a contrast between 0.2 and 0.31, the participant saw the manipulated grating two times and had a mixed percept one time. This means that the midpoint of those contrasts (0.255) was assigned a dominance level of 2.5/11 = 0.23. Then, for the trials between 0.21 and 0.32, the participant saw the manipulated grating three times, giving the middle contrast (0.265) a dominance level of 3/11 = 0.28.

### Bayesian hierarchical model

To analyse the effect of manipulated sensory evidence on dominance, we constructed a Bayesian hierarchical model^1^. Details of the model specification, its inference and Bayesian model comparison are described in the Supplementary Material. With this model we simultaneously estimated the participant-level psychometric response curve parameters, as well as their noise levels and group-level response curve parameters, for both the congruent and the incongruent imagery condition. This allowed us to (1) test whether there was in general an effect of imagery and an interaction with sensory evidence; (2) establish whether those effects were present for each individual participant, and finally; (3) estimate the size and variability of an effect, both at the group-level as well as per participant.

The result of the inference procedure was the posterior distribution of both group-level and participant-level parameters, for each condition. In addition, it provided the distribution over the *differences* in these parameters between conditions. Using Bayesian model comparison, we tested whether these differences differed from zero. For example, consider the parameter *u* that represents the bias in the psychometric curve (i.e. its horizontal offset; see Fig. 1). Let *δ*_*u*_ = *u*^congruent^ − *u*^incongruent^ be the difference in this parameter between the two conditions. We write 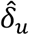 to indicate the group-level difference and 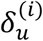 to indicate the difference for parameter *μ* for participant *i*. Using the Savage-Dickey method (Wagenmakers, 2010), we computed the Bayes factor that quantifies how much more likely the observations are under the alternative model *H*_+_ in which there is a difference between the conditions, versus under the null model *H*_0_ of no difference. That is, we obtain:

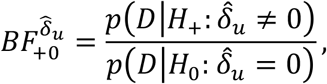

for the test of a difference in *u* at the group-level. Note that all these tests were performed on the one posterior distribution of the full hierarchical model. This caused participant-level parameters to be pulled towards the group-mean (a phenomenon known as *shrinkage*), which is the Bayesian hierarchical way of mitigating false alarm rates (Kruschke, 2018). To interpret the (logarithm of the) Bayes factors, we used the interpretation table provided by Lee and Wagenmakers (2014), as presented in Table 1.

**Table 1.**
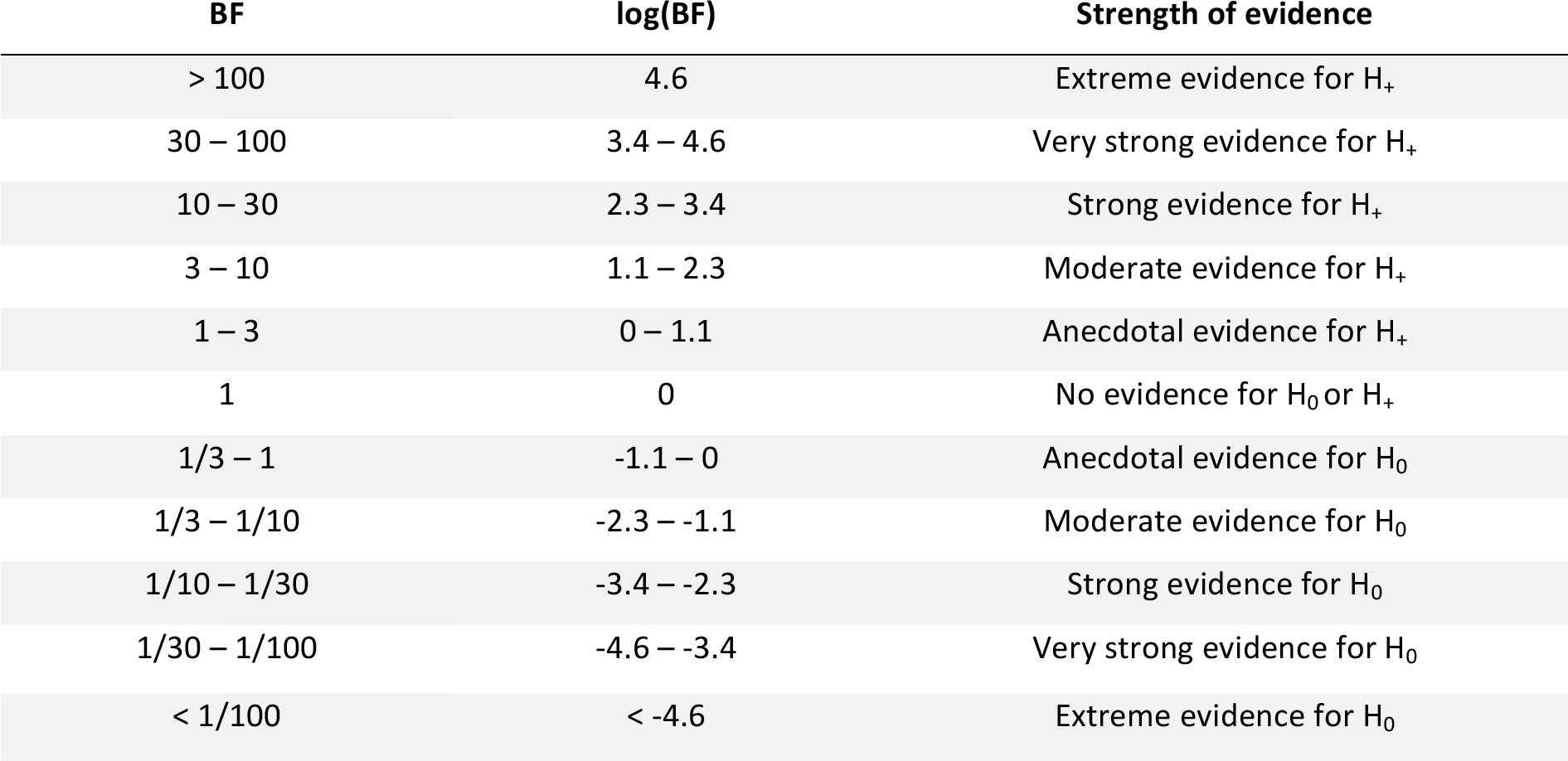
Interpretation of Bayes factors (BF). Interpretation of Bayes factors according to Jeffreys (1961) and Lee and Wagenmakers (2014). First column shows the BF which shows how much more likely the alternative hypothesis (H_+_) is compared to the null hypothesis (H_0_). The second column shows the log of the BF, which is used for visualization in Figure 3 and 4. The third column shows the interpretation of the BFs.

## Results

### Replication imagery priming group-effect

First, we aimed to replicate the main imagery priming effect reported in earlier studies. In these studies, the imagery effect was determined by first removing all mixed trials and then assessing whether participants perceived the imagined percept more often than chance (Bergmann, Genç, Kohler, Singer, & Pearson, 2016; Keogh & Pearson, 2011; Pearson et al., 2011; Pearson et al., 2008a). A two-sided one-sample t-test revealed that the proportion of trials in which participants perceived the imagined percept was indeed significantly larger than 0.5 (*M* = 0.53, *SD* = 0.05, *t*(58) = 3.78, *p* = 0.0004, *d* = 0.6). Note that for the analyses reported below, we did not remove the mixed trials, but instead used them as providing information about the relationship between sensory evidence and dominance (see section *Deriving dominance per contrast*).

### Response bias

To assess whether there was a response bias, we looked at the responses to mock rivalry trials. During these trials, a mock rivalry stimulus was presented, which should elicit the response of a mixed percept. The response bias was assessed by determining whether participants were more likely to indicate that they had perceived the imagined percept during mock trials. Even though there was still an indication of mock priming in some participants, there was no correlation between mock priming and binocular priming (*r* = 0.03, *p* = 0.81; *BF* = 0.17). This means that people who had a strong response bias did not show a strong imagery effect on the binocular rivalry trials. Therefore, the imagery effects could not be explained by response bias.

### Main effect of imagery

For the main analyses, we used a hierarchical Bayesian model to fit psychometric curves on the dominance responses during congruent and incongruent imagery. The main effect of imagery is reflected in the difference in *bias* (Fig. 1) between the two conditions. A positive difference indicates that the bias during congruent imagery was smaller than during incongruent imagery, so that less contrast was needed to reach the same dominance level. Thus, a positive difference indicates a *priming* effect of imagery. A negative difference shows that during congruent imagery more contrast was needed compared to incongruent imagery to achieve the same contrast levels, indicating an *adaptation* effect of imagery. The posterior estimates of the differences, together with their corresponding log Bayes factors are shown in Fig. 3.

**Figure 3.**
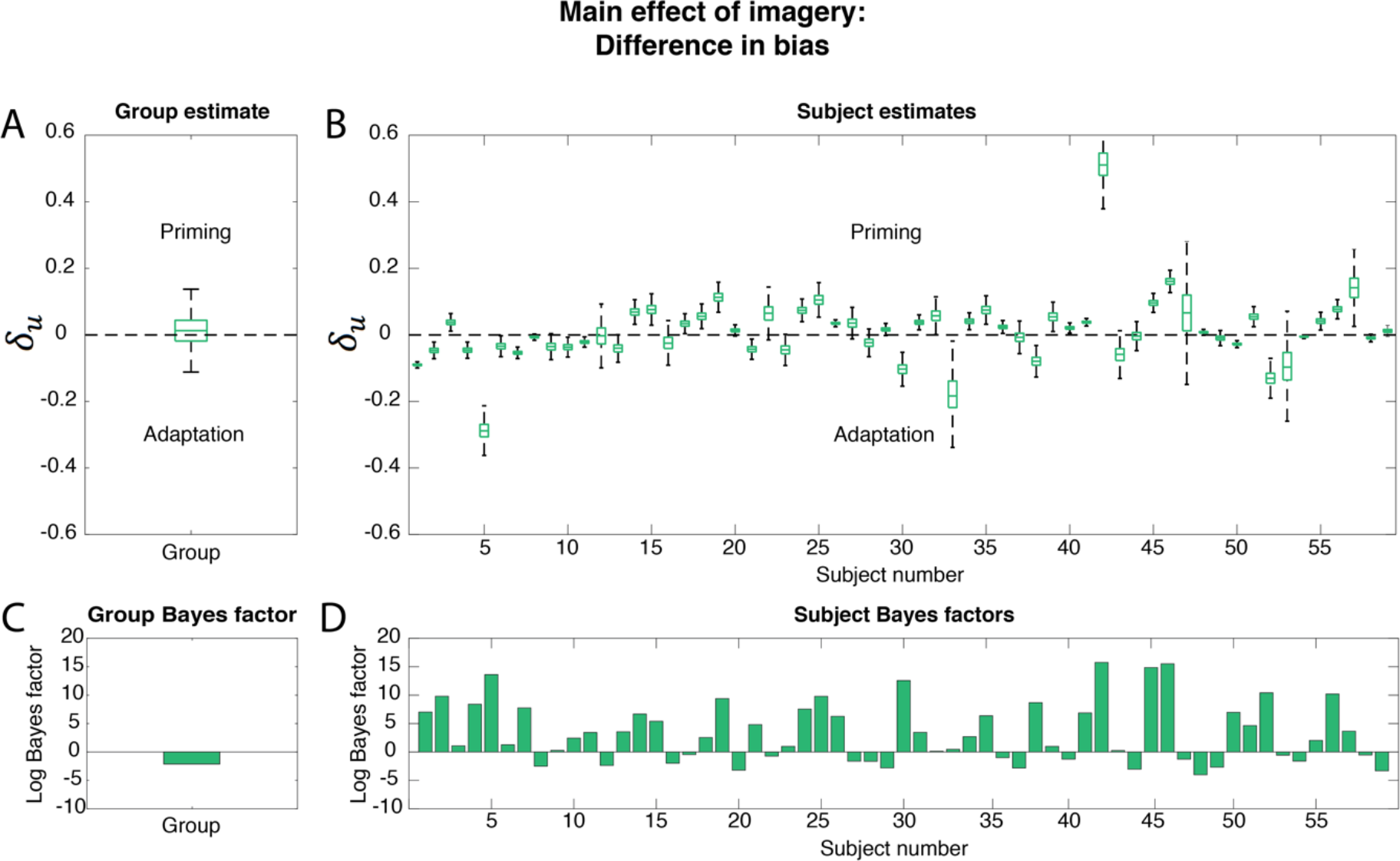
Main effect of imagery: difference in bias. The difference in bias 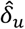 between incongruent and congruent imagery was tested. A positive difference reflects a higher bias for incongruent imagery which indicates a priming effect while a negative difference indicates an adaptation effect. Boxplots reflect the distribution of the posterior samples. Outliers are not shown. The bottom of the box represents the first quantile, the middle line represents the median and the top, the third quantile. The distance between the bottom and the top of the box reflects the variance of the samples and is a measure of the uncertainty of the estimate. (A) Posterior group estimate of the difference. (B) Posterior subject estimates of the difference; each boxplot represents one participant. The logs of the BFs for the comparison between H_+_ (difference) and H_0_ (no difference) are shown. A positive log(BF) indicates evidence in favour of H_+_ and a negative log(BF) indicates evidence in favour of H_0_. (C) Group log(BF) and (D) subject log(BF); each bar represents one subject. The order of the participants is the same in (B) and (D).

The group log(BF) is −2.14 (Fig. 3C), which is interpreted as moderate evidence for the absence of an effect (Table 1). Note that this is in contrast to the results of the t-test reported above which showed a significant group priming effect. However, individual subject level Bayes factors were very high with a median log(BF) of 2.44, indicating strong evidence for the presence of an effect (Fig. 3D). The reason for the discrepancy between the group-level effect and the participant-level effects is the inconsistency of the effect across participants. That is, the direction of the effect varied strongly between participants (Fig. 3B), leading to a null effect on the group level (Fig. 3A). A large proportion of participants showed a strong priming effect associated with a high Bayes factor (Fig. 4A; upper right quadrant). However, almost as many participants showed a reliable adaptation effect (Fig. 4A; upper left quadrant). Furthermore, there was also a proportion of participants who reliably did not show an effect, i.e. for which there was evidence in favour of H_0_ (Fig. 4A; log(BFs) below 0). See Fig. 6 for examples of psychometric curve fits for a few different participants.

**Figure 4.**
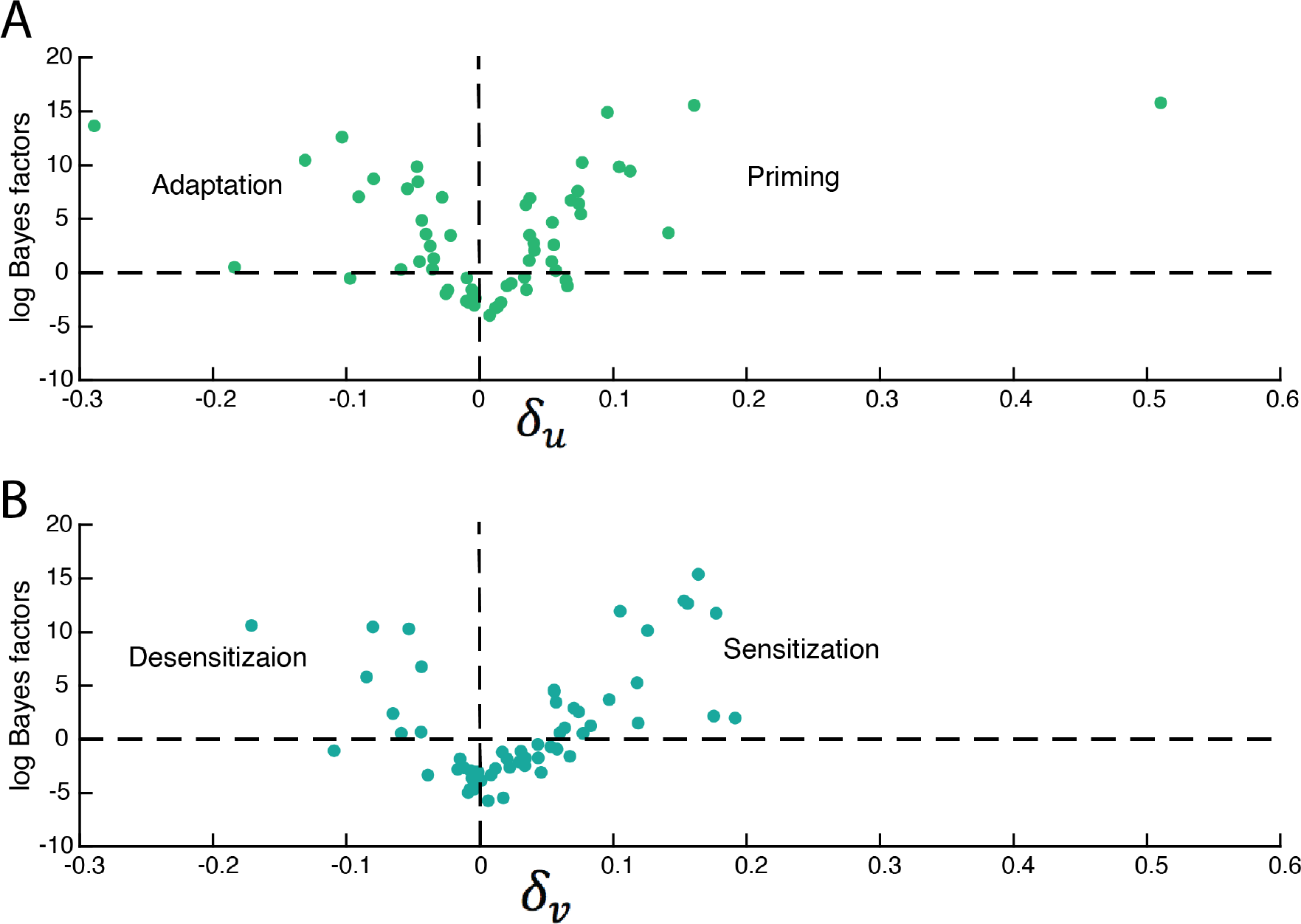
Direction and Bayes factor of effects per participant. On the x-axis, the median of the posterior samples for the difference between the two conditions is plotted. On the y-axis, the log(BF) of the effect is plotted. Negative log(BFs) indicate evidence in favour of H_0_ and positive log(BFs) indicate evidence in favour of H_+_. Dots represent individual participants. (A) Main effect of imagery reflected in the difference in bias 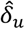 between conditions. (B) Interaction between imagery and perception reflected in the difference in discrimination sensitivity 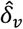 between the two conditions.

### Interaction with sensory evidence

The interaction between imagery and sensory evidence is reflected in a difference in slope or *discrimination sensitivity* (see Fig. 1) between congruent and incongruent imagery. The discrimination sensitivity reflects how much increase in contrast is needed to achieve an increase in dominance. A low slope means that small changes in contrast are enough to influence perception whereas a high slope means that larger changes are necessary. Thus, a positive difference in discrimination sensitivity means that the slope is higher for incongruent imagery, indicating that congruent imagery *increases* sensitivity to bottom-up sensory input. In contrast, a negative difference indicates that imagery *decreases* sensitivity. The results are shown in Fig. 5.

**Figure 5.**
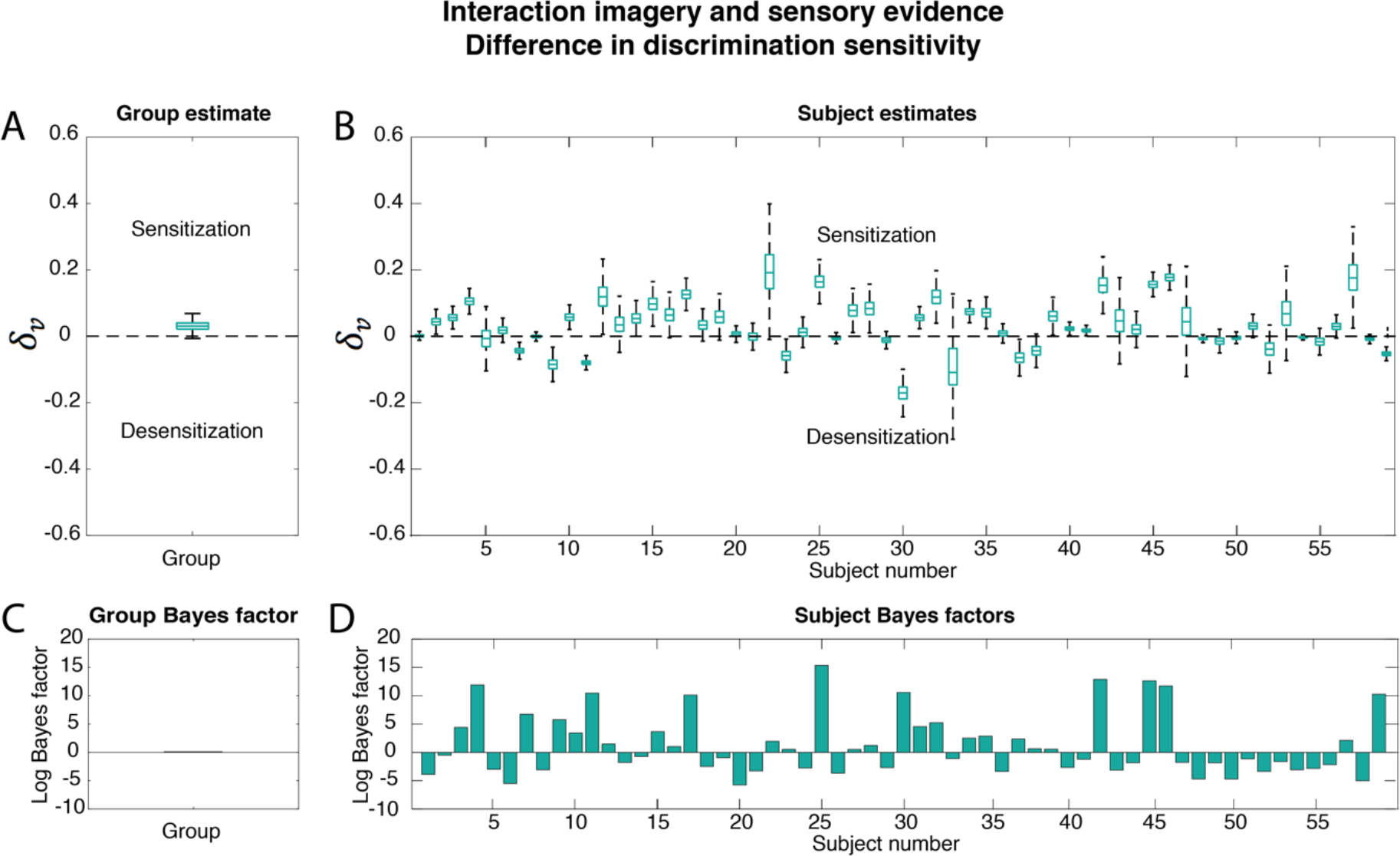
Interaction between imagery and sensory evidence: difference in discrimination sensitivity. The difference in discrimination sensitivity 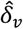 between incongruent and congruent imagery was tested. A positive difference reflects an increase in sensitivity for congruent imagery whereas a negative difference indicates a decrease in sensitivity. Boxplots reflect the distribution of the posterior samples as in Fig. 3 (A) Posterior group estimate of the difference. (B) Posterior subject estimates of the difference, each boxplot represents one participant. The logs of the BFs for the comparison between H_+_difference_ / H_0_no_difference_ are shown. (C) Group log(BF) and (D) subject log(BFs), each bar represents one subject. The order of the participants is the same in (B) and (D) and the same as in Fig. 3.

The group BF is 0.02, which reflects no clear evidence for either H_+_ or H_0_ (Fig. 4A & 4C). The individual participant’s BFs for the difference in *discrimination sensitivity* (Fig. 5D) are less high than those for the difference in *bias* (Fig. 3D). Most participants do not show a clear interaction with sensory evidence (Fig. 4B; log(BFs) below 0). However, there are still a number of participants who do show a reliable increase in discrimination sensitivity for congruent imagery (Fig. 4B; upper right quadrant, e.g. Fig. 6A), and participants who show a reliable decrease (Fig. 4B; upper left quadrant).

**Figure 6.**
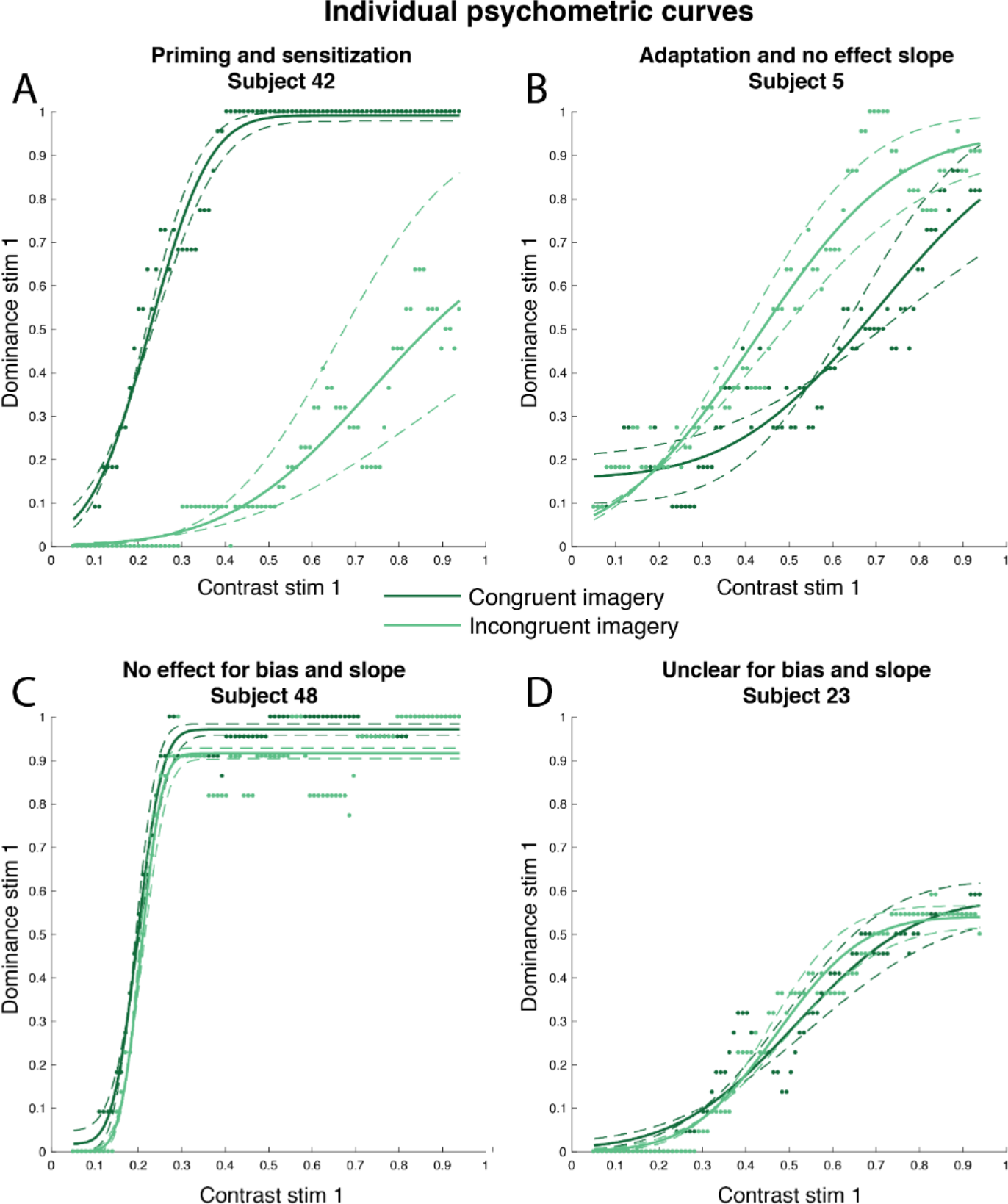
Psychometric curves for four participants. The dominance for the manipulated stimulus is shown on the y-axis and its contrast on the x-axis. Dots represents individual trials, solid lines the posterior estimate of the fitted curve and dashed lines its uncertainty. Dark green shows dominancy during congruent imagery and light green during incongruent imagery. (A) A participant showing a reliable imagery priming effect as well as an increase in sensitivity during congruent imagery. (B) A participant showing a reliable imagery adaptation effect and no difference in discrimination sensitivity between conditions. (C) A participant reliably showing no main imagery effect and no interaction with sensory evidence. (D) A participant whose data did not allow dissociation between H_+_ and H_0_.

To test the robustness of our results, we checked whether the Bayesian model gave similar results if we used a wider or narrower prior (see Supplementary Material). Changing the prior did not significantly change the results, indicating that the findings were not influenced strongly by the choice of prior parameters.

### Relation between vividness and imagery effects

We also investigated the relation between subjective vividness ratings and imagery’s influence on conscious perception. We first aimed to replicate the findings from earlier studies. To this end, we binned the trials into four levels based on the vividness ratings and for each bin we calculated the priming effect, operationalized as the proportion of non-mixed trials that the imagined percept was perceived, as in Pearson et al. (2011). In line with earlier findings (Pearson, Rademaker, & Tong, 2011), there was a main effect of vividness on imagery priming (*F*(2.45,134.72) = 3.77, *p* = 0.018, Huyn-Feldt-corrected for violation of sphericity). This effect was linear (*F*(1,55) = 6.29, *p* = 0.015), indicating that the effect of imagery on binocular rivalry was stronger for more vivid imagery. In contrast to earlier findings, we did not find a correlation between VVIQ and imagery priming (*r* = 0.09, *p* = 0.47, *BF* = 0.17), showing that off-line VVIQ scores did not predict participant’s imagery priming.

Finally, we tested whether the difference in psychometric curve parameters derived by using all data (including mixed trials) showed a relationship with imagery vividness. Since it was impossible to obtain binned psychometric curve values, we only focused on group-level correlations with VVIQ and averaged vividness ratings. There was no correlation between the difference in bias and VVIQ (*r* = −0.03, *p* = 0.80, *BF* = 0.17) or averaged vividness rating (*r* = −0.06, *p* = 0.66, *BF* = 0.18). There was also no correlation between the difference in discrimination sensitivity and VVIQ (*r* = −0.02, *p* = 0.87, *BF* = 0.17) or averaged vividness rating (*r* = −0.04, *p* = 0.76, *BF* = 0.17). This indicates that the individual differences in the interaction between imagery and perception reported above are not easily linked to self-report measurements of imagery vividness.

## Discussion

In this study, we set out to characterize how imagery influences conscious perception. We tested whether the influence of imagery was influenced by bottom-up sensory input and to what extent it differed between participants. We first replicated the priming group-effect of imagery on binocular rivalry. Using standard analyses, the results indicated that participants are more likely to perceive a stimulus that they had just imagined. However, using a hierarchical Bayesian model to estimate full psychometric response curves per participant revealed a variety of effects. First, whereas a large proportion of participants did indeed show an imagery priming effect, almost as large a group showed an imagery adaptation effect and another large proportion reliably showed no effect. Furthermore, while most participants did not show a clear interaction between imagery and sensory evidence, some participants still showed a reliable increase in sensitivity for congruent imagery and some participants even showed a clear decrease in sensitivity. These results indicate that the influence of imagery on conscious perception is more complicated than previously reported and that there are large individual differences in the nature of this process.

The existence of both priming and adaptation effects of imagery is in line with the fact that both of these effects are found for prior perception (Knapen et al., 2007; Pearson et al., 2008). Presenting a stimulus prior to a binocular rivalry display can bias dominance towards or away from the presented stimulus depending on the contrast. Brief, low-contrast stimuli tend to result in a priming effect whereas long, high-contrast stimuli result in an adaptation effect. The idea is that prior perception pre-activates low-level neural populations involved in later stimulus processing during binocular rivalry. This prior activation builds up depending on the duration and contrast of the stimulus, leading to a stronger priming effect until it reaches some threshold after which it leads to adaptation (Knapen et al., 2007). Given the large neural overlap between imagery and perception, it is not surprising that imagery can also elicit both priming and adaptation effects. If the same mechanism is at work, we would expect that imagery adaptation is caused by stronger and/or longer low-level activation than imagery priming. We did not find a relationship between self-report measurements of imagery vividness and the direction of this effect. This might be due to the fact that vividness does not fully capture the strength and the duration of imagery. Future research should investigate whether there is a direct link between neural activation and imagery’s influence on conscious perception.

Furthermore, most participants did not show an effect of imagery on discrimination sensitivity. However, for some participants there was a clear effect, indicating that the strength of the influence of imagery can depend on the bottom-up sensory evidence. This means that the influence of imagery on perception might not always be linear. An increase in discrimination sensitivity indicates that imagery enhances differences in sensory input and can thereby facilitate detecting sensory input congruent to our internal goals. In contrast, a decrease in discrimination sensitivity means that the visual system becomes less sensitive to sensory input congruent to imagery. Participants showing desensitization usually showed a main adaptation effect, suggesting that desensitization might also be caused by some form of neural fatigue caused by strong activations.

An obvious next step is to investigate the prevalence of these different effects in the population. One straightforward way to do this is to extent the Bayesian model put forward here to include a term that assigns participants to a group depending on their effects (primer, adapter, sensitizer, etc.) and test this model on new data. An important question that can then be answered is to what extent these individual differences relate to psychopathology. For example, people with very strong priming effects were more likely to perceive sensory input congruent with their imagery even when incongruent signals were stronger. Perceiving internally generated content despite counteracting sensory evidence has been put forward as an explanation for hallucinations (Powers, Kelley, & Corlett, 2016). In contrast, underweighting internal signals with respect to external signals has been put forward as an explanation for perceptual differences in autism spectrum disorder (Skewes, Jegindø, & Gebauer, 2015). Investigating to what extent these effects correlate with other perceptual and cognitive variations could have important clinical implications.

We were unable to estimate the full psychometric curve for some participants because they either needed very low contrast values to reach a 100% dominance (e.g. Fig. 6C), or too high values (e.g. Fig. 6D). This is due to large individual differences in the strength of eye dominance (Mapp, Ono, & Barbeito, 2003). This means that for some people we were unable to fully characterize the effects. In order to circumvent this issue, future research could use different techniques to assess eye dominance (Ding, Naber, Gayet, Van der Stigchel, & Paffen, 2018) or determine the stimulus strength needed to estimate the full psychometric curve in a separate session. It is likely that this will increase the number of participants that show an effect.

In conclusion, our results show that imagery influences conscious perception differently for different people. Imagery can bias perception towards and away from the imagined stimulus and can increase and decrease sensitivity to congruent sensory input. These interactions are probably supported by a neural mechanism involving imagery’s recruitment of low-level visual processes. Future research should investigate whether these individual differences correspond to clinical variations. This study furthermore highlights the importance of investigating within-participant effects and proposes a way to do so.

## Acknowledgements

SEB and MG were supported by VIDI grant number 639.072.513 of The Netherlands Organization for Scientific Research (NOW). ND was supported by institutional funding. MH was supported by the European Research Council Consolidator grant number 647209 and the European Research Council Advanced grant number 743086. The authors would like to thank Surya Gayet for his extremely useful comments on the design of the experiment, Rebecca Keogh for important input regarding the binocular rivalry imagery task and Lynn Le for her invaluable role in data collection.

## Footnotes

1 A discussion of the advantages of Bayesian inference is beyond the scope of this paper, but we refer the interested reader to (Berger & Berry, 1988; Wagenmakers, 2007).

