## Supplementary Material for "Individual differences in the influence of mental imagery on conscious perception"

### 1 Power calculation

We performed a power-calculation using G-power (Erdfelder et al., 2009) to determine the required number of participants needed to investigate individual differences. Previous research investigating individual differences in imagerys effect of binocular rivalry have reported a range of correlation strengths: 0.38 (Bergmann et al., 2015), 0.57 (Keogh and Pearson, 2014), 0.45 (Keogh et al., 2016) and 0.73 (Pearson et al., 2011). Assuming a correlation of 0.35, a power of 0.8, an alpha of 0.05 and a two-tailed test, the power calculation showed that we required a sample size of 59.

### 2 Bayesian hierarchical model

#### 2.1 Model definition

The model is based on the psychometric curve by Wichmann and Hill (2001). It defines the response function as

$$f(x; g, l, u, v) = g + (1 - g - l)\Phi\left(\frac{x - u}{v}\right) \quad , \tag{1}$$

in which $x$ is the manipulated contrast, $g \in [0, 1]$ is the *guess rate*, $l \in [0, 1]$ the *lapse rate*, $u \in [0, 1]$ the *bias*, $v \in \mathbb{R}$ the *discrimination sensitivity* and $\Phi(\cdot)$ the cumulative Normal distribution. For examples of such curves, we defer to the main document. From this definition we construct a Bayesian hierarchical model as follows. For participant $i = 1, \ldots, N$, condition $k \in \{1, 2\}$ and observed stimulus contrast level $j = 1, \ldots, M$ we define the likelihood

$$\begin{aligned} \mu_{ijk} \mid g_{ik}, l_{ik}, u_{ik}, v_{ik}, c_j &= f(c_j; g_{ik}, l_{ik}, u_{ik}, v_{ik}) \\ d_{ijk} &\sim \text{Normal}\left(\mu_{ijk}, \tau_i\right) \quad . \end{aligned} \tag{2}$$

Here, the parameter $\tau_i$ is the precision of subject $i$, corresponding to the inverse of its noise level. This term is estimated by placing a vague prior distribution over it, that is

$$\tau_i \mid \alpha_\tau, \beta_\tau \sim \text{Gamma}\left(\alpha_\tau, \beta_\tau\right) \tag{3}$$

with $\alpha_\tau = \beta_\tau = 0.001$.

Hierarchy is introduced to the model in a standard fashion by centering the prior distribution for each participant-level parameter on the corresponding population-level parameter. That is,

$$\begin{aligned} g_{ik} \mid \hat{g}_k, \tau_g &\sim \text{TNormal}_0^1\left(\hat{g}, \tau_g\right) \\ l_{ik} \mid \hat{g}_k, \tau_l &\sim \text{TNormal}_0^1\left(\hat{l}, \tau_l\right) \\ u_{ik} \mid \hat{u}_k, \tau_u &\sim \text{TNormal}_0^1\left(\hat{u}, \tau_u\right) \\ v_{ik} \mid \hat{v}_k, \tau_v &\sim \text{Gamma}\left(\hat{v}^2\tau_v, \hat{v}\tau_v\right) \quad , \end{aligned} \tag{4}$$

where $\text{TNormal}_a^b(\mu, \tau)$ is the Normal distribution with mean $\mu$ and precision $\tau$, but with its domain truncated to the interval $[a, b]$.

For the population-level parameters (indicated with the ˆ symbol) we define (hyper-)priors in which we encode our informed prior beliefs on the psychometric curve. We define

$$\begin{aligned} \hat{g}_k \mid \mu_{\hat{g}}, \tau_{\hat{g}} &\sim \text{TNormal}_0^1(\mu_{\hat{g}}, \tau_{\hat{g}}) \\ \hat{l}_k \mid \mu_{\hat{l}}, \tau_{\hat{l}} &\sim \text{TNormal}_0^1(\mu_{\hat{l}}, \tau_{\hat{l}}) \\ \hat{u}_k \mid \mu_{\hat{u}}, \tau_{\hat{u}} &\sim \text{TNormal}_0^1(\mu_{\hat{u}}, \tau_{\hat{u}}) \\ \hat{v}_k \mid \mu_{\hat{v}}, \tau_{\hat{v}} &\sim \text{Gamma}(\mu_{\hat{v}}^2 \tau_{\hat{v}}, \mu_{\hat{v}} \tau_{\hat{v}}) \ , \end{aligned} \tag{5}$$

where we set $\mu_{\hat{g}} = 0.02$, $\tau_{\hat{g}} = \tau_{\hat{l}} = \tau_{\hat{v}} = 1/0.1^2$, $\mu_{\hat{l}} = \mu_{\hat{v}} = 0.1$, $\mu_{\hat{u}} = 0.5$ and $\tau_{\hat{u}} = 1/0.5^2$, representing the expected group-level curves.[1]. We set the prior on the guess rate higher than the lapse rate because it is more difficult to reach 100 percent dominance while the other stimulus is fixed at 0.4 contrast than it is to reach 0 percent dominance (for more details, see main text).

Similarly, we estimate the precision parameters that determine the amount of variability of participants around the group-level means. As we have no strong prior beliefs about this variability, we use vague priors (Lee and Vanpaemel, 2018), defined via

$$\begin{aligned} \tau_g \mid \alpha_{\tau_g}, \beta_{\tau_g} &\sim \text{Gamma}(\alpha_{\tau_g}, \beta_{\tau_g}) \\ \tau_l \mid \alpha_{\tau_l}, \beta_{\tau_l} &\sim \text{Gamma}(\alpha_{\tau_l}, \beta_{\tau_l}) \\ \tau_u \mid \alpha_{\tau_u}, \beta_{\tau_u} &\sim \text{Gamma}(\alpha_{\tau_u}, \beta_{\tau_u}) \\ \tau_v \mid \alpha_{\tau_v}, \beta_{\tau_v} &\sim \text{Gamma}(\alpha_{\tau_v}, \beta_{\tau_v}) \ , \end{aligned} \tag{6}$$

where we set $\alpha_m = \beta_m = 0.001$ for $m \in \{\tau_g, \tau_l, \tau_u, \tau_v, \tau_i\}$.

### 2.2 Inference

Inference of the model is done via Gibbs Markov chain Monte Carlo sampling, using the JAGS software (Plummer, 2003). We computed 4 parallel chains of 100 000 samples and verified convergence of the sampler visually as well as via computing the potential scale reduction factor $\hat{R}$ for each parameter, and asserting that its value remained below the common heuristic of 1.1 (Brooks and Gelman, 1997). This procedure results in the approximated posterior distribution $p(\boldsymbol{\theta} \mid D, c, \boldsymbol{\psi})$ where $\boldsymbol{\theta}$ contains all latent parameters, $D \in [0,1]^{M \times N \times K}$ is the observed indicated dominance, $c \in [0,1]^M$ is the manipulated contrast and $\boldsymbol{\psi}$ is the set of hyperparameters of the model.

### 2.3 Model comparison using Bayes factors

The approximated posterior distribution of the latent parameters implies a distribution over the differences of these parameters. Consider for example the group-level bias term. We have available the posterior distribution of $\hat{u}_k$ for both conditions, so we can derive the implied posterior distribution over $\hat{\delta}_u = \hat{u}_2 - \hat{u}_1$. Subsequently, we test whether $\hat{\delta}_u \neq 0$, that is we compare the alternative hypothesis $H_+ : \hat{\delta}_u \neq 0$ with the null model in which there is no effect, that is, $H_0 : \hat{\delta}_u \neq 0$.

As $H_0$ is a special case of the full model, we can use the Savage-Dickey method to conveniently compute the corresponding Bayes factors for these tests (Dickey, 1971; Wagenmakers et al., 2010). For this method, we approximate both the posterior as well as the prior model (that is, $p(\theta \mid D, c, \boldsymbol{\psi})$ and $p(\theta \mid c, \boldsymbol{\psi})$), and compute the ratio of these densities at $\hat{\delta}_u = 0$, i.e.:

$$BF_{+0}^{\hat{\delta}_u} = \frac{p(D \mid H_+, \boldsymbol{\psi})}{p(D \mid H_0, \boldsymbol{\psi})} = \frac{p(\hat{\delta}_u = 0 \mid H_+, c, \boldsymbol{\psi})}{p(\hat{\delta}_u = 0 \mid H_+, D, c, \boldsymbol{\psi})} \ . \tag{7}$$

The Bayes factors for the different comparisons, both at the group-level and the participant-level, are shown in the main text.

### 2.4 Robustness check

To check that our results were not too influenced by our choices of hyperparameter settings, we ran the analysis again, once with broader priors and once with narrower priors. The results are shown

[1]Note that it is typically easier to think of standard deviation and variance rather than precision. Hence we transform our beliefs about standard deviation into beliefs about precision via the identity $\tau = 1/\sigma^2$.

in Figure S1. The pattern of Bayes factors using a wide or narrow prior is very similar to that observed using the original prior. The correlations between the log(BFs) is shown in Figure S2. All correlations are higher than 0.94, indicating that differences in the prior did not substantially change our findings.

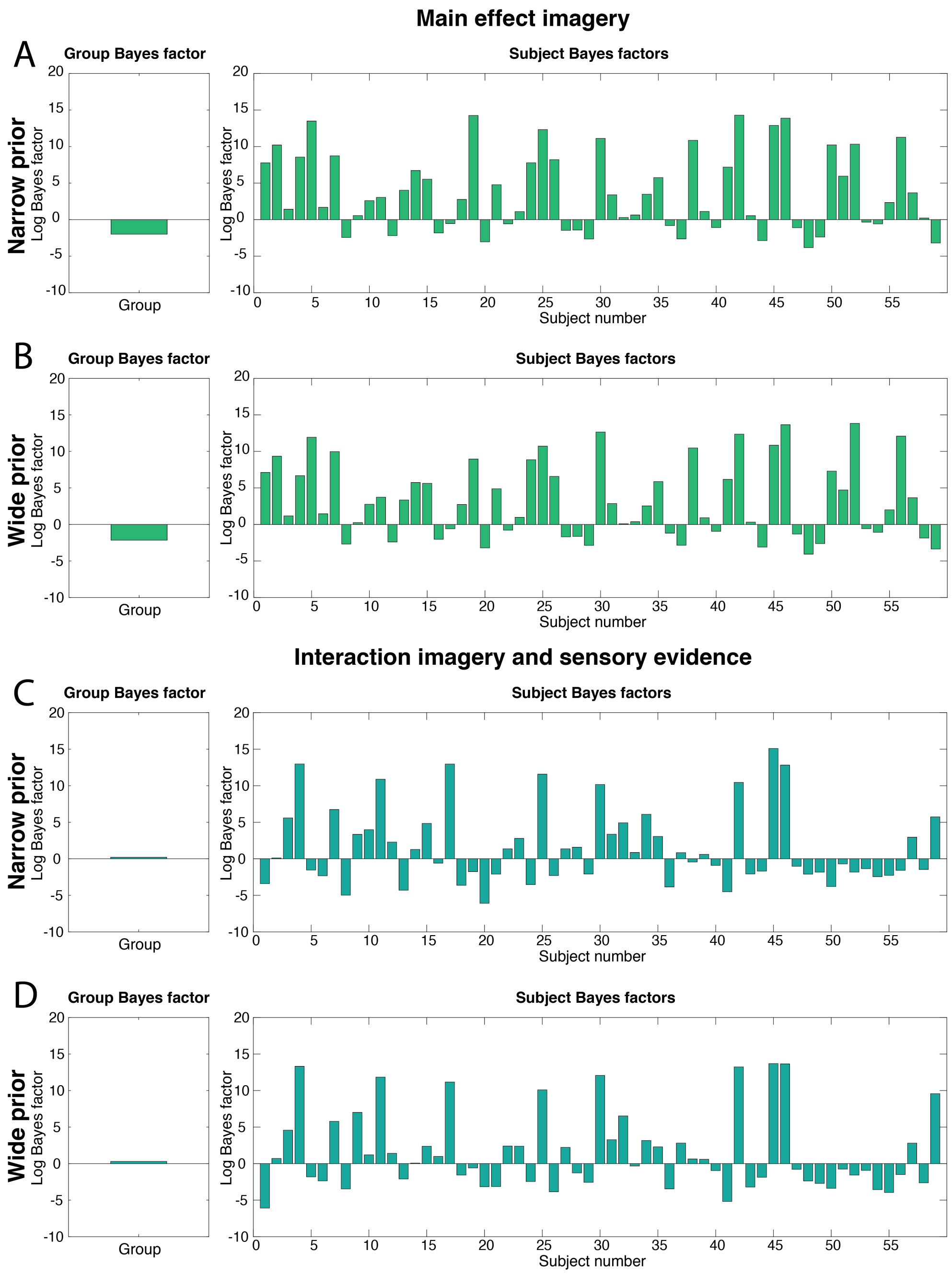


Figure S1: **Bayes factors robustness check: Bayes factors.** Bayes factors with narrow priors and with wide priors. (A) BFs for the difference in bias with narrow priors. (B) BFs for the difference in bias with wide priors. (C) BFs for the difference in discrimination sensitivity with narrow priors. (D) BFs for the difference in discrimination sensitivity with wide priors.

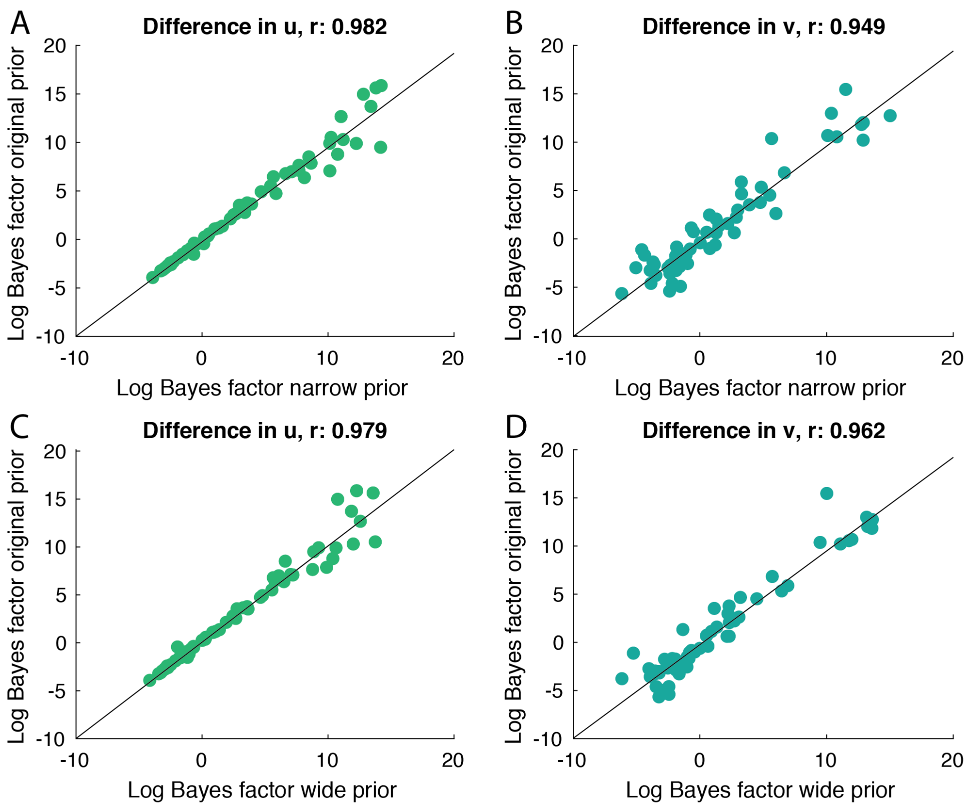


Figure S2: **Bayes factors robustness check: correlations.** Comparison between BFs using the original priors reported in the main text and using narrower (A, C) or wider (B, D) priors. Each circle represents a participant.
